# A Humanized Animal Model Predicts Clonal Evolution and Therapeutic Vulnerabilities in Myeloproliferative Neoplasms

**DOI:** 10.1101/2020.11.12.378810

**Authors:** Hamza Celik, Ethan Krug, Christine RC Zhang, Wentao Han, Nancy Issa, Won Kyun Koh, Hassan Bjeije, Ostap Kukhar, Maggie Allen, Tindao Li, Daniel AC Fisher, Jared Fowles, Terrence N. Wong, Matthew C. Stubbs, Holly K. Koblish, Stephen T. Oh, Grant A. Challen

## Abstract

Myeloproliferative neoplasms (MPNs) are chronic blood diseases with significant morbidity and mortality. While sequencing studies have elucidated the genetic mutations that drive these diseases, MPNs remain largely incurable with a significant proportion of patients progressing to rapidly fatal secondary acute myeloid leukemia (sAML). Therapeutic discovery has been hampered by the inability of genetically-engineered mouse models to generate key human pathologies such as bone marrow fibrosis. To circumvent these limitations, here we present a humanized animal model of myelofibrosis (MF) patient-derived xenografts (PDXs). These PDXs robustly engrafted patient cells which recapitulated the patient’s genetic hierarchy and pathologies such as reticulin fibrosis and propagation of MPN-initiating stem cells. The model can select for engraftment of rare leukemic subclones to identify MF patients at-risk for sAML transformation, and can be used as a platform for genetic target validation and therapeutic discovery. We present a novel but generalizable model to study human MPN biology.

**STATEMENT OF SIGNIFICANCE:** Although the genetic events driving myeloproliferative neoplasms (MPNs) are well-defined, therapeutic discovery has been hampered by the inability of murine models to replicate key patient pathologies. Here, we present a patient-derived xenograft (PDX) system to model human myelofibrosis that reproduces human pathologies and is amenable to genetic and pharmacological manipulation.

## INTRODUCTION

Myeloproliferative neoplasms (MPN) are clonal hematologic diseases characterized by the aberrant proliferation of one or more myeloid lineages, and progressive bone marrow (BM) fibrosis. BCR-ABL negative MPNs are characterized by an excess of red blood cells [polycythemia vera (PV)] or platelets [essential thrombocythemia (ET)], or by the deposition of reticulin fibers in the BM [myelofibrosis (MF)](1). MF is the deadliest MPN subtype with median survival for patients of approximately five years, and can occur as a *de novo* neoplasm or progression from pre-existing ET or PV. The most commonly mutated genes in MPNs are *JAK2* (2–4), *MPL* (5) and *CALR* (6,7), which are responsible for disease initiation. The acquisition of additional mutations in epigenetic regulators, transcription factors, and signaling components can modify the course of the disease and contribute to disease progression and leukemic transformation (8,9). *JAK2, MPL*, and *CALR* mutations share a hallmark of aberrant JAK-STAT activation. As such, the JAK1/2 inhibitor ruxolitinib is a front line therapy (in particular for MF) and can alleviate constitutional symptoms of the disease (10), but does not eliminate the malignant clone and has minimal impact on BM fibrosis and overall survival (11), highlighting the need for more effective therapies.

As the genetic basis of MPN has been elucidated, murine models have been developed to dissect disease origins and mechanisms. Transgenic and conditional knock-in mice have shown that physiological expression of *JAK2*^V617F^ alone can induce a PV- or ET-like MPN *in vivo* (12), and the hematopoietic stem cell (HSC) pool contains the disease-initiating potential (13,14). However, current mouse models do not recapitulate the clinical heterogeneity, genetic composition, or morphological features of MPN. Most notably, these models do not typically generate robust reticulin fibrosis in the BM, the most significant MF pathology. The lack of faithful pre-clinical MPN models presents a major barrier to developing effective therapies.

To circumvent the limitations of conventional mouse models, patient-derived xenografts (PDXs) are increasingly used as tools for pre-clinical drug development and target validation, and have been shown to recapitulate the pathology and genetics of the patients from which they are derived in various malignancies (15). Technological developments in immunodeficient mouse strains (16,17) have enabled successful xenotransplantation of normal human CD34+ hematopoietic stem and progenitor cells (HSPCs), as well as patient samples from myelodysplastic syndromes (MDS) and acute myeloid leukemia (AML) subtypes that were previously refractory to engraftment. However, efforts to generate MF PDXs have been largely unsuccessful due to poor engraftment potential of MPN-disease initiating cells (18), or require specialized techniques (19).

In a previous study, we demonstrated that genetic inhibition of the polycomb repressive 2 (PRC2) co-factor JARID2 in MPN patient samples permitted engraftment in NSGS mice (20). This caused us to revisit methodology to determine if technological advances might facilitate a generalizable MPN PDX system. Here, we present a generalizable method for xenotransplantation of human MF CD34+ cells into newer generation immunodeficient mice by X-ray guided intra-tibial injection, which results in robust engraftment. These PDXs reproduce the hallmarks of MF including splenomegaly, myeloid maturational arrest and, most importantly, reticulin fibrosis in the BM. Genetic analysis shows faithful maintenance of MF patient clonal architecture within the engrafted cell population. Enhancing the utility of the model, we show this system can predict for clonal progression to secondary AML (sAML) in patients samples obtained far prior to clinical manifestation, and can be manipulated to provide a platform for genetic and pharmacological target validation in MF. Thus, this model presents a robust and malleable platform to study MPN genetics and therapeutics.

## RESULTS

### Credentialing a Humanized Animal Model of Myelofibrosis

In our previous study of tumor suppressors in chronic myeloid neoplasms, genetic inhibition of the PRC2 co-factor *JARID2* by lentiviral shRNA permitted sustained engraftment of CD34+ cells from MF patients in the blood, bone marrow and spleens of NSGS mice accompanied by marked reticulin fibrosis (20). However, we noted evidence of engraftment and modest reticulin fibrosis in recipients of MF patient cells transduced with the negative control shRNA vector. This encouraged us to revisit the methodology to determine if techniques could be adapted for a generalizable PDX model of MF. CD34+ cells were freshly isolated from the peripheral blood mononuclear cell (PBMC) fraction from MF patients (**Supplementary Fig. 1**). Selection criteria for patient samples included blast count <5%, with over 80% of patient samples having no detectable blasts. MF CD34+ cells were cultured for 16-hours in serum free media with hematopoietic cytokines (SCF, FLT3L and TPO), then transplanted into sublethally (200 rads) irradiated NSGS mice via X-ray guided intra-tibial (IT) injection (**Fig. 1A**). NSGS mice express humanized SCF, GM-CSF and IL3 (16) and demonstrate improved myeloid engraftment from normal human CD34+ hematopoietic stem and progenitor cells (HSPCs) and AML subtypes that were previously difficult to engraft (21,22). The technical adaptation of IT injection has facilitated PDX models from difficult to engraft diseases such as CMML (23). Comparison of IT versus intravenous injection showed superior engraftment for both normal and neoplastic human CD34+ cells (**Fig. 1B**). We reasoned these technological advances may be sufficient to generate reliable MF PDXs.

**Figure 1.**
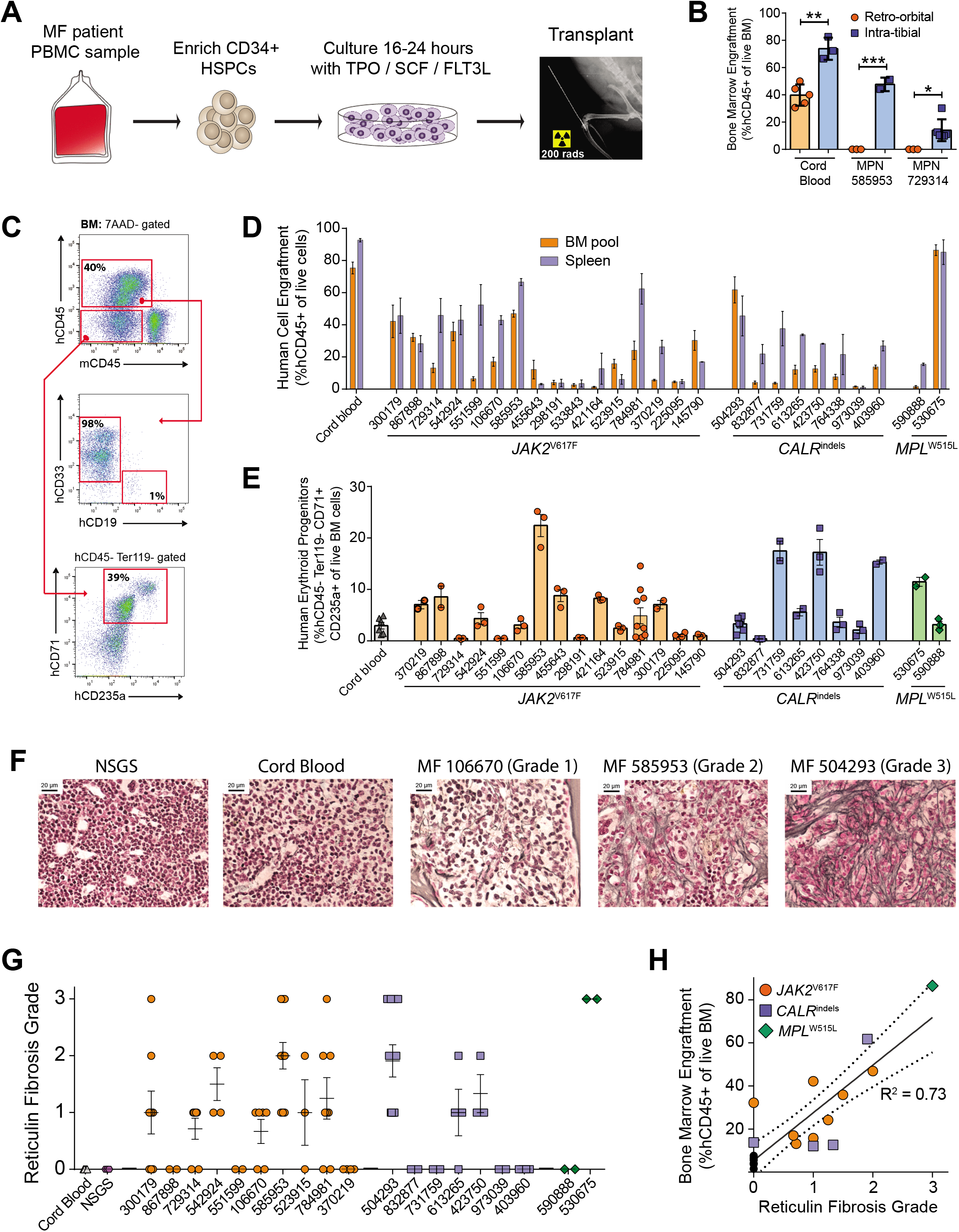
Credentialing a Humanized Animal Model of Myelofibrosis. **A**, X-ray guided intra-tibial injection of CD34+ cells from MF patients. **B**, Comparison of BM engraftment 12-weeks post-transplant resulting from either retro-orbital or intra-tibial injection of CD34+ cells from cord blood and MF patient samples. **C**, Flow cytometric identification of engrafted human cells in NSGS mice. **D**, Engraftment levels of MF patient cells in BM and spleens of NSGS mice 12-weeks post-transplant. **E**, Engraftment of erythroid progenitor cells (hCD45-Ter119-CD71+ CD235a+) of MF patient cells in NSGS BM 12-weeks post-transplant. **F**, Reticulin staining showing fibrosis in the BM of NSGS mice transplanted with MF patient samples, but not in untransplanted mice or mice transplanted with cord blood CD34+ cells. Representative reticulin fibrosis grading is indicated. **G**, Quantification of degree of reticulin fibrosis from each patient sample. **H**, Correlation of reticulin fibrosis grade with BM engraftment. Dashed lines represent 95% confidence intervals. Error bars indicate mean ± S.E.M. * *p*<0.05, ** *p*<0.01, *** *p*<0.001.

While the engraftment of human cells in the peripheral blood was highly variable between patient samples (**Supplementary Fig. 2A**), virtually all NSGS mice that received >100,000 CD34+ MF patient cells showed robust engraftment in the BM (**Fig. 1C**) and spleen at 12-week post-transplant (**Fig. 1D**). Consistent with MPN, engrafted human cells were almost exclusively CD33+ myeloid cells (**Fig. 1C**) or immature erythroid progenitors (**Fig. 1E**). There was no difference in BM engraftment between the injected and contralateral legs (**Supplementary Fig. 2B**). Engraftment levels were not correlated with number of CD34+ cells transplanted (**Supplementary Fig. 2C**), but rather inherent properties of individual samples. Analysis revealed other pathologies characteristic of MPN such as activated JAK/STAT signaling (**Supplementary Fig. 2D**) and increased blood counts (**Supplementary Fig. 2E,F**). The most aggressive patient samples were able to induce a lethal MPN in NSGS mice (**Supplementary Fig. 2G**). Most strikingly, most MF PDXs showed marked reticulin fibrosis in the BM and spleen compared to mice transplanted with cord blood CD34+ cells, or NSGS mice that were irradiated and then monitored for 12-weeks (**Fig. 1F**). Grading of reticulin fibrosis severity (**Fig. 1G**) showed that the degree of fibrosis correlated with overall engraftment levels of MF cells in the BM (**Fig. 1H**). Thus, this methodology provides a robust and reproducible model for establishing PDXs from MF patient samples.

### MF HSCs Show Robust Engraftment in NSGS Mice

To ensure the engrafted MF cells in NSGS mice retained the molecular properties of the patient cells, single cell RNA-seq was performed. PBMCs from two independent cord blood and MF patient samples were compared to the engrafted hCD45+ cells from the BM of NSGS mice 12-weeks post-transplant. UMAP analysis showed that the PBMCs from both cord blood samples clustered together, but the MF PBMCs clustered distinctly from the normal cells as well as from each other (**Fig. 2A**). Although *JAK2*^V617F^ was the driver mutation in both these MF patients, the separation from each other was likely driven by the different cooperating mutations (e.g. MF 585953 = *TET2, ASXL1;* MF 784981 = *SETBP1, CUX1, ZRSR2*). But importantly, the independent cell clusters were maintained in engrafted PDX cells. hCD45+ cells isolated from the BM of NSGS mice 12-weeks post-transplanted grouped in the same UMAP clusters as the PBMCs from the donors pre-transplant (**Fig. 2A**). Annotation of cell clusters using marker gene expression showed that the MPN-specific cell clusters were enriched for myeloid cells, platelets, and HSPCs in both the patients and the PDXs (**Fig. 2A**). Gene set enrichment analysis (GSEA) revealed that significantly upregulated gene expression in the neoplastic cells was enriched cytokine signaling pathways in both the patient and PDX cells (**Fig. 2B**), a hallmark of MPN (24). This informs that the distinct molecular profiles of MF patient cells are maintained in this PDX model.

**Figure 2.**
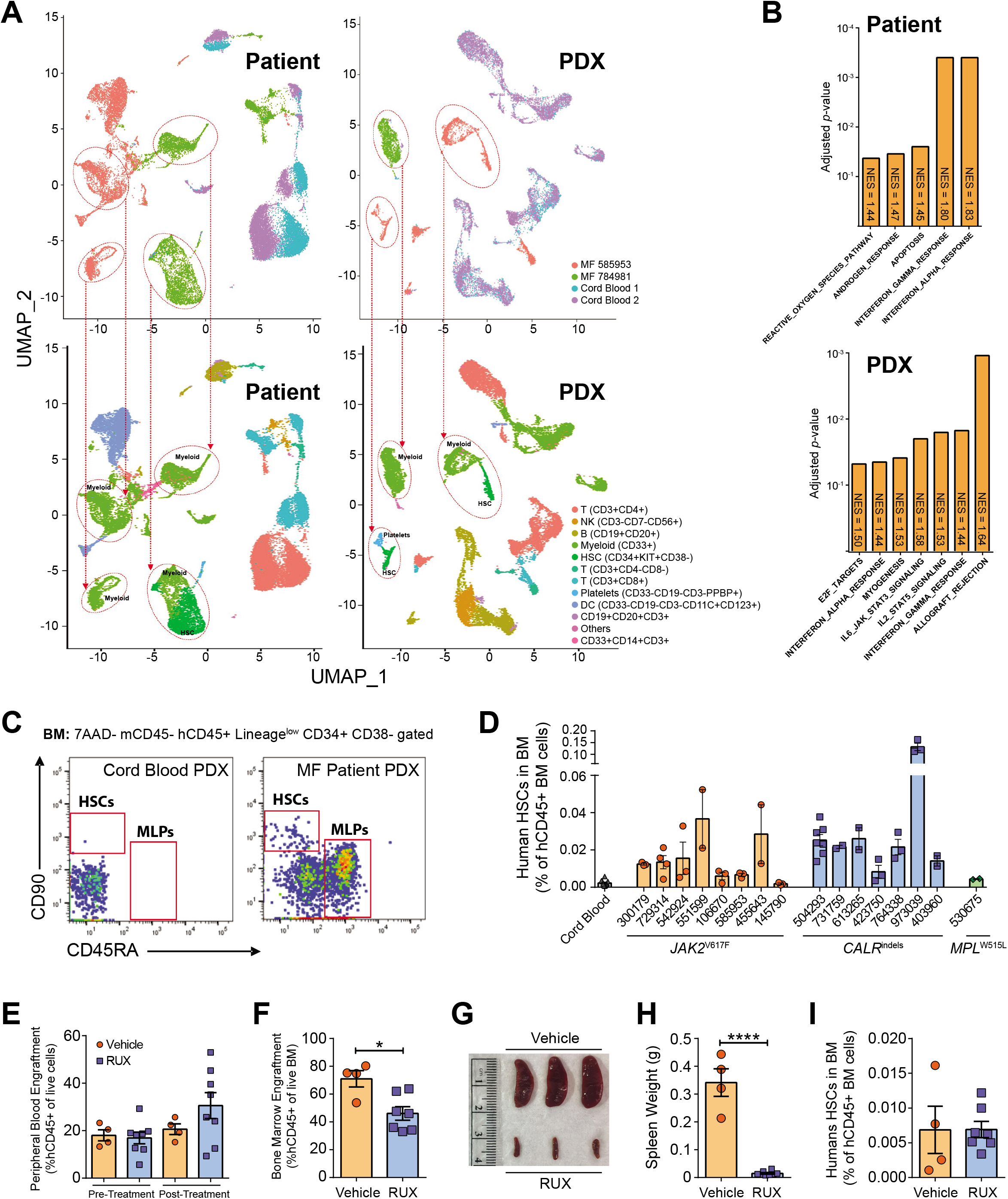
MF HSCs Show Robust Engraftment in NSGS Mice. **A**, UMAP plots of single cell RNA-seq data showing clustering of PBMC cells from two cord blood samples and two MF patients (Patients) and hCD45+ cells from the same donors isolated from the BM of NSGS mice 12-weeks post-transplant (PDX). The top panels show clustering of samples by genotype in Patients and PDX. The bottom panels show identification of cell populations by marker genes. Dashed red lines indicate population identities of MPN-only cell clusters. **B**, Gene set enrichment analysis showing pathways significantly upregulated in MPN patient cells in the Patients (top) and PDX (bottom). **C**, Flow cytometric identification of human HSCs (hCD45+ Lineage^low^ CD34+ CD38-CD45RA-CD90+) gated in the BM of NSGS mice. **D**, Quantification of human HSCs in the BM of NSGS mice 12-weeks post-transplant. **E**, Four-week peripheral blood engraftment of MF 504293 patient cells prior to treatment initiation, and four-weeks after ruxolitinib therapy. **F**, BM engraftment of MF patient cells four-weeks post-treatment. **G**, Representative spleen images from mice of the different treatment groups. **H**, Spleen weights of NSGS four-weeks post-treatment. **H**, Quantification of human HSC abundance in the BM of NSGS mice post-treatment. Error bars indicate mean ± S.E.M. * *p*<0.05, **** *p*<0.0001.

Experimental models implicate HSCs as the disease-initiating cell population in MPN (13,14). Moreover, MPN founding mutations such as *JAK2*^V617F^ and *CALR* indels can be detected in humans with clonal hematopoiesis (CH)(25–27), further suggesting a HSC origin as the mutational reservoir. Most MF therapies are likely ineffective because they fail to eradicate this mutant HSC population in patients. We sought to determine if the MPN mutant HSCs were propagated in our PDX model, which could provide a system to evaluate the impact of novel therapeutics specifically on the disease-initiating cell population. Flow cytometric analysis of the BM of recipient NSGS mice showed that human HSCs (Lineage^low^ CD34+ CD38-CD45RA-CD90+) were not maintained in animals xenografted with cord blood CD34+ cells (**Fig. 2C**). This is likely due to the inflammatory environment driving HSCs towards terminal differentiation as opposed to selfrenewal (28). Conversely, analysis of NSGS mice xenografted with CD34+ cells from MF patients showed that human HSCs were readily detectable in most cases (**Fig. 2D**), suggesting that this is a permissive niche for the propagation of MPN disease-initiating cells. To determine if MPN mutant HSCs could be serially transplanted, 1×10^6^ hCD45+ cells from primary PDXs were transferred to secondary NSGS recipients using the same technical approach. Despite robust engraftment in primary transplantation (**Supplementary Fig. 2H**), MF patient cells were unable to engraft or propagate disease pathologies to secondary recipients (**Supplementary Fig. 2I**). This however does indicate that the samples initially transplanted, while neoplastic, were not transformed and the resultant engraftment was not due to leukemic subclones.

Given that MPN disease-initiating HSCs were propagated, we sought to benchmark the therapeutic relevance of the system by evaluating the effect of ruxolitinib as current standard of care for MF. NSGS mice were xenografted with MF CD34+ cells, and then four-weeks posttransplant were randomized for treatment with ruxolitinib (or vehicle control) based on engraftment of human cells in the peripheral blood. Engraftment and pathology of MF patient cells were evaluated after four-weeks of therapy. Treatment of MF cells with ruxolitinib in NSGS mice recapitulated the patient experience. While ruxolitinib treatment did not affect the burden of MF cells in the peripheral blood (**Fig. 2E**), burden of human cells was significantly reduced in the BM (**Fig. 2F**) accompanied by a dramatic reduction in splenomegaly (**Fig. 2G,H**). But consistent with human patients, treatment with ruxolitinib did not affect the abundance of human HSCs in the BM (**Fig. 2I**).

### MF Patient Clonal Architecture is Maintained in PDX Models

The genetic hierarchy of MPN is hypothesized to influence treatment response to JAK inhibitors. To credential this model as an avatar for pre-clinical screening, exome sequencing was performed on MF patient cells prior to transplantation and hCD45+ cells 12-weeks posttransplant to determine if the patient’s clonal architecture was preserved in PDXs (**Supplementary Table 1**). The mutational landscape observed in MF patients was exquisitely maintained in the xenografted cells from NSGS mice (**Supplementary Fig. 3**). There was high reproducibility for the same patient sample xenografted into different NSGS mice (**Fig. 3A**) and the variant allele fraction (VAF) of driver mutations showed excellent correlation between patient and PDX cells (**Fig. 3B**). In almost all PDXs, the engrafted human cells were virtually exclusively of the myeloid lineage (CD33+). However, xenografted CD34+ cells from MF patient 300179 was the only sample to generate significant lymphoid progeny in NSGS mice (**Fig. 3C**), which allowed comparison of mutant allele burden in different cell lineages and analysis of the developmental trajectory of this patient. In this patient, heterozygous *DNMT3A^P307L^* and *TET2*^P1962L^ mutations were found in all cells from the patient donor sample and the xenografted cells. Homozygous *JAK2*^V617F^ and *TP53*^M246V^ mutations were detected at a lower VAF in the bulk post-transplant hCD45+ cells compared to the donor sample, due to the fact these variants were completely absent from the B-cells (CD19+) in the PDX (**Fig. 3C**). This suggests the B-cells were the progeny of a non-neoplastic CH clone with *DNMT3A*^P307L^ and *TET2*^P1962L^ mutations. For MPN evolution, this clone subsequently acquired homozygous *JAK2*^V617F^ and *TP53*^M246V^ mutations either in a HSC or downstream myeloid progenitor cell which drove excess myeloid differentiation and clinical manifestation (**Fig. 3D**).

**Figure 3.**
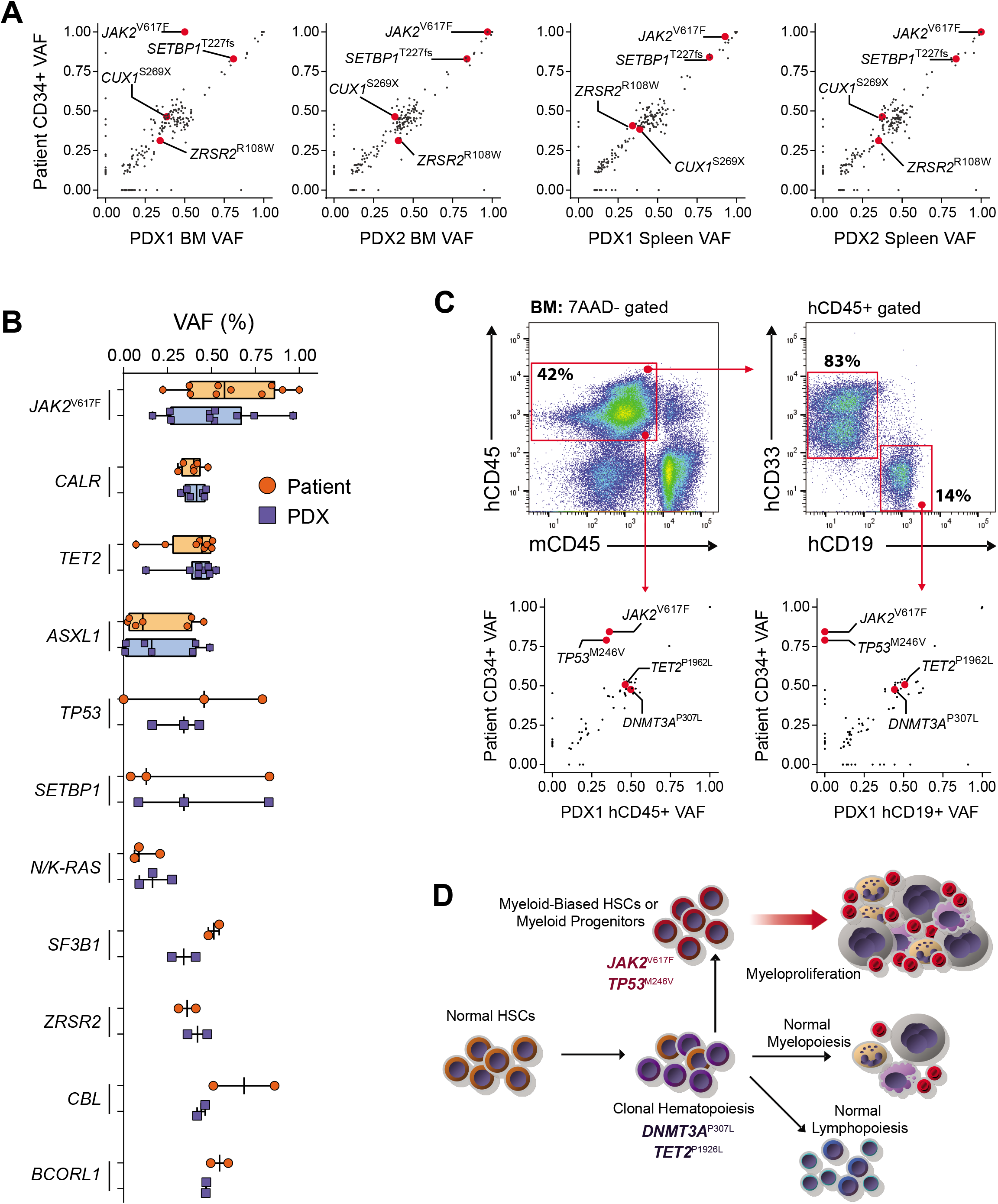
MF Patient Clonal Architecture is Maintained in PDX Models. **A**, Comparison of the variant allele fraction (VAF) of mutations in CD34+ cells from MF patient 784981 versus hCD45+ cells isolated from BM and spleens of NSGS mice 12-weeks posttransplant. **B**, Comparison of VAFs of recurrently mutated genes in this cohort between MF patient CD34+ cells and PDX-derived hCD45+ cells. **C**, Flow cytometry plots showing isolation of different cell populations from PDX of MF patient 300179 and the VAF of mutations in each respective cell populations. **D**, Model for clonal evolution in MF patient 300179 based on mutational profile of cell populations derived from PDX.

### PDX System can Predict Clonal Evolution to Secondary AML

One of the most feared complications of MPN is disease progression to sAML. Up to 20% of patients with MF transform sAML, which is driven by the acquisition of additional co-operating mutations (29). Patients with post-MF sAML have a dismal prognosis, with a median survival of only six-months (9). Earlier identification of MF patients who are more susceptible to sAML transformation could stratify such individuals for alternative interventions. Such predictions are not possible with conventional genomic approaches. For the PDXs from patient MF 504293, exome sequencing detected a pathogenic *EZH2*^Y663H^ mutation in the xenografted human cells which were not identified in the input CD34+ cells directly isolated from the patient (**Fig. 4A**). Review of patient history uncovered that this individual eventually underwent disease progression to sAML (**Fig. 4B**). Strikingly, clinical sequencing performed at the time of sAML diagnosis identified the same *EZH2*^Y663H^ mutation present in the PDX, which had never been observed in previous clinical sequencing performed throughout the patient history (**Fig. 4C**). Of note, the PBMC sample used for PDX input was obtained from this patient 516 days prior to sAML diagnosis. This implies that the selection pressure of xenotransplantation facilitated expansion of a very rare subclone (below the level of detection of conventional exome sequencing) in the PDX model which was responsible for leukemic transformation. As the xenografted cells were obtained from the patient in the chronic MF stage years before sAML diagnosis (**Fig. 4B,C**), this suggests that this PDX system might serve as a prediction model for patients with rare, rising sAML subclones who are at-risk for leukemic progression. To examine this, droplet digital PCR (ddPCR) was performed with an *EZH2*^Y663H^-specific probe on an independent patient sample banked at the same time as the specimen used for xenotransplantation. ddPCR analysis (**Supplementary Fig. 4A**) indeed confirmed the presence of very rare *EZH2*^Y663H^-mutant cells in the PBMCs of this patient (0.17% of cells), with enrichment in the CD34+ cell population (0.63% of cells), although still far below the sensitivity of detection of conventional exome sequencing (**Fig. 4D**). These data suggest that rare sAML subclones are detectable in chronic MF patients years before clinical manifestation, which are readily identifiable by the PDX model.

**Figure 4.**
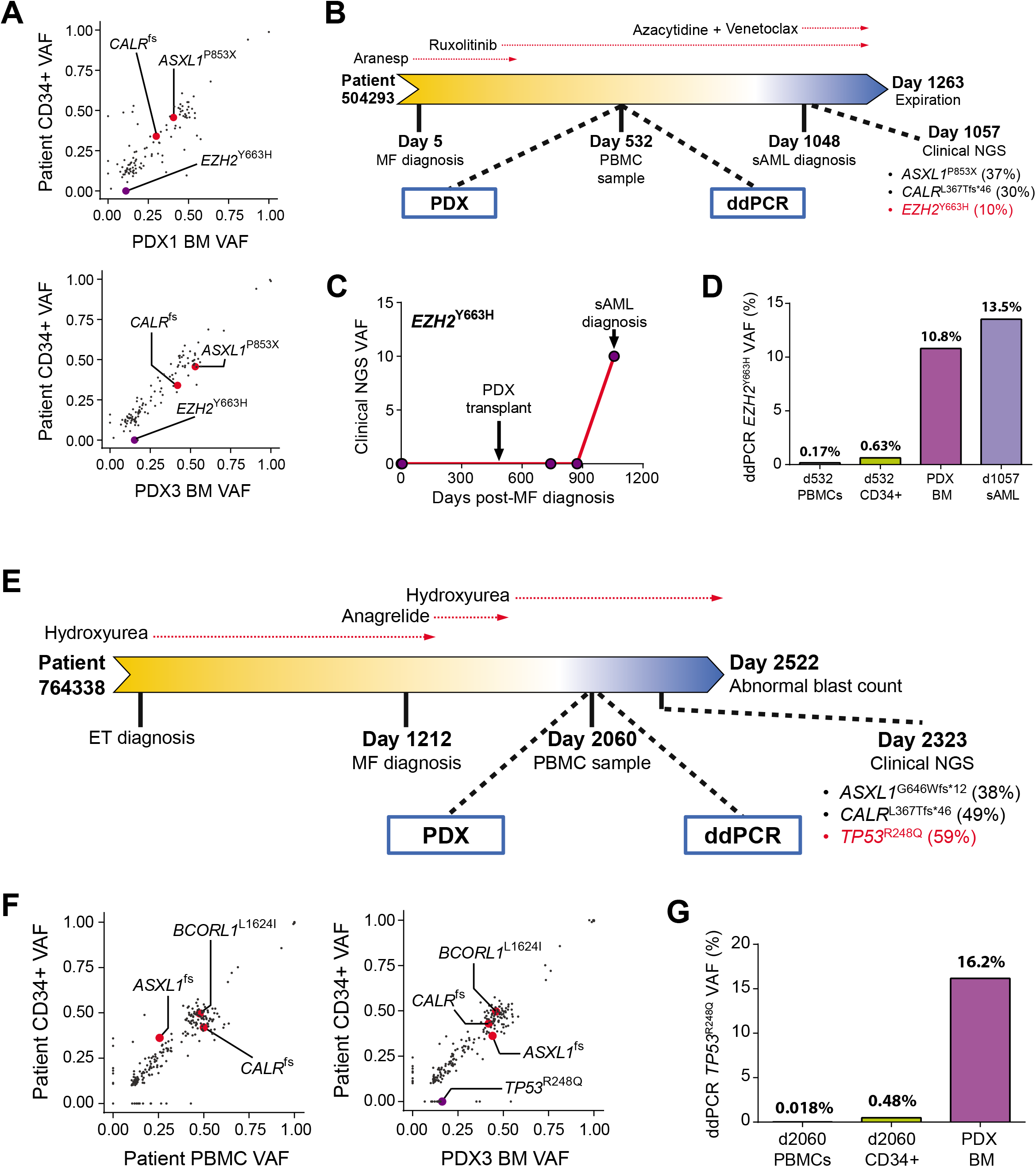
PDX System can Predict Clonal Evolution to Secondary AML. **A**, Comparison of the variant allele fraction (VAF) of mutations in CD34+ cells from MF patient 504293 versus hCD45+ cells isolated from BM of NSGS mice 12-weeks post-transplant. **B**, Clinical course of MF patient 504293. **C**, VAF of *EZH2*^Y663H^ variant from clinical NGS during clinical course of MF patient 504293. **D**, Quantification of *EZH2*^Y663H^ variant from indicated samples by ddPCR. **E**, Clinical course of MF patient 764338. **F**, Comparison of VAF of mutations in CD34+ cells from MF patient 764338 versus PBMCSs and hCD45+ cells isolated from BM of NSGS mice 12-weeks post-transplant. **G**, Quantification of *TP53^R248Q^* variant from indicated samples by ddPCR.

Analysis of other exome sequencing cases identified another individual (MF 764338) in which a pathogenic variant was detected exclusively in the human cells from the PDX, but not in the input CD34+ cells at time of collection. At the most recent biopsy, this patient was flagged for having an abnormal blast count and is currently under consideration for a bone marrow transplant. Immediately prior to this diagnosis, clinical sequencing identified the *TP53^R248Q^* mutation in the patient PBMCs, which was the first time in the patient history this variant had been observed (**Fig. 4E**). The hCD45+ cells from the PDX were found to harbor the *TP53*^R248Q^ mutation, one of the most common *TP53* variant in de novo AML (30) and post-MPN sAML (31), at a VAF of 15-20% (**Fig. 4F**). We returned to an independent PBMC sample from the time of collection which was used for xenotransplantation to determine if *TP53*^R248Q^-mutant could be identified by ddPCR. This analysis showed that in whole PBMCs, *TP53*^R248Q^-mutant cells were at the level of sensitivity of ddPCR (**Supplementary Fig. 4B**). But in CD34+ cells enriched from this timepoint, a distinct population of *TP53*^R248Q^-mutant cells (0.48% of cells) could be identified (**Fig. 4G**). Cumulatively, these two cases demonstrate that rare disease-transforming clones are present in MF patients long prior to diagnosis of clinical progression. While these clones are undetectable using current clinical sequencing methodology, they can be readily identified in this PDX model.

### Genetic and Pharmacological Validation of Novel Therapeutic Targets in MF

We sought to establish the utility of this model for genetic and pharmacological target validation. Proof-of-principle focused on PIM kinase inhibition. The PIM family (PIM1-3) of kinases regulate a number of cellular processes through phosphorylation of a variety of target substrates. The proto-oncogenic potential of PIM kinases was initially described in MYC-driven lymphomagenesis (32), and PIM inhibition (PIMi) is being explored for a variety of solid tumors (33,34) as well as hematopoietic malignancies (35,36). In *JAK2*-mutant MPN cell lines, PIMi has been shown restrain cell proliferation and overcome drug resistance through destabilization of MYC and suppression of pro-survival signaling pathways (37). PIM1 kinase is also upregulated in MF CD34+ cells regardless of their genetic background (38). Resultantly, PIMi has been presented as a therapeutic approach in *JAK2*-mutant MPN in combination with JAK/STAT inhibition. Our system provides a unique opportunity to validate these *in vitro* studies with cell lines using primary patient samples *in vivo*.

To determine the specificity of PIMi, cord blood (normal control) or MF patient CD34+ cells were targeted for *PIM1* deletion via CRISPR/Cas9 RNP nucelofection (39). A gRNA targeting the inert *AAVS1* locus was used as a CRISPR/Cas9 negative control. Analysis of targeted cells via deep-sequencing 72-hours post-nucleofection confirmed high efficiency editing (**Supplementary Fig. 5A**). Edited CD34+ cells were transplanted into NSGS recipients as described, and mice were sacrificed 12-weeks post-transplant to evaluate function of targeted cells. While loss of *PIM1* had no effect on the engraftment of cord blood-derived cells, *PIM1*-targeted MF cells showed markedly compromised engraftment in the blood (**Supplementary Fig. 5B**), reduced splenomegaly (**Supplementary Fig. 5C**) and reduced burden in the bone marrow (**Supplementary Fig. 5D**) which was not due to absence of targeted *PIM1* populations (**Supplementary Fig. 5A**). These data suggest that *PIM1* may be selectively required for MF cell function.

To evaluate pharmacological potential, PDXs were established from MF patient samples and then randomized for treatment with the pan-PIMi INCB053914 (40), either as a single agent or in combination with the JAK/STAT inhibitor ruxolitinib. Treatment was initiated four-weeks posttransplant, then effects were evaluated after four-weeks of treatment (**Fig. 5A**). In comparison to vehicle-treated control mice, treatment with either INCB053914 or ruxolitinib alone modestly inhibited engraftment of MF cells and splenomegaly for each of three individual patient samples to varying degrees. However, combination therapy markedly inhibited engraftment of patient cells (**Fig. 5B**) and reduced peripheral blood counts (**Supplementary Fig. 5E**). Strikingly, while treatment with either single agent had no effect on the burden of human HSCs in the BM, combination therapy was effective in eradicated the *JAK2*^V617F^-mutant MPN-initiating cell population (**Fig. 5C**). As a dysregulated cytokine profile is a feature of MPN patients as well as mouse MPN models which may drive disease progression of the neoplastic clone, the treatment effect on serum cytokine levels was evaluated. While single agent therapy reduced pro-inflammatory cytokine levels to varying degrees, combination therapy had potent antiinflammatory effects, exemplified by the effects on IL-1b, IL-6 and IL-8 (**Supplementary Fig. 5F**). These results show that PIMi in combination with JAK/STAT inhibition specifically inhibits MPN, particularly *JAK2*^V617F^-mutant cells, and present this combination as an attractive therapeutic approach for MF patients.

**Figure 5.**
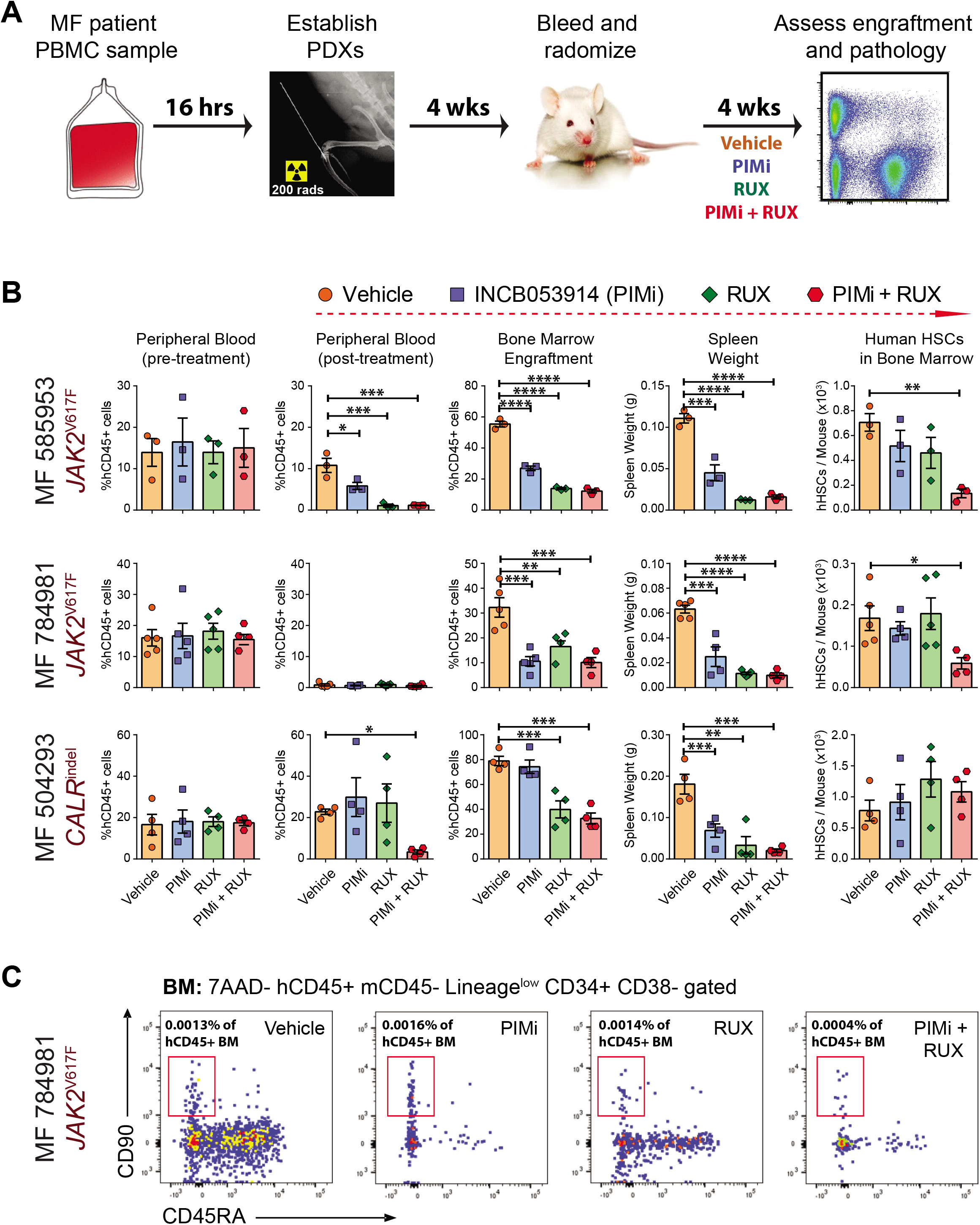
Genetic and Pharmacological Validation of Novel Therapeutic Targets in MF. **A**, Schematic for MF combination therapy using PDX model. **B**, Cumulative data showing MF patient cell peripheral blood engraftment pre-treatment, and post-treatment blood and BM engraftment, spleen weight and human HSC burden in the BM. **C**, Representative flow cytometry plots showing human HSC burden from NSGS mice transplanted with MF 784981 and then exposed to the indicated treatments. Error bars indicate mean ± S.E.M. * *p*<0.05, ** *p*<0.01, *** *p*<0.001, **** *p*<0.0001.

## DISCUSSION

MPNs are largely incurable hematopoietic diseases. While symptoms can be managed in many patients, patients remain at risk for thrombohemorrhagic complications and disease transformation. Curative therapies would alleviate a substantial burden on public health, but require methods to effectively eradicate the founding MPN clone. New therapeutic approaches based on patient molecular profiles are needed, which has been hampered by the inability of genetically engineered mouse models to recapitulate key MF pathologies such as reticulin fibrosis. Here, we present a system for establishing PDXs, which accurately propagates the genotypes and phenotypes of MF patients. The model produces robust engraftment of patient-derived cells in all major hematopoietic organs, sustains disease-initiating MPN HSCs, and disseminates the most critical MF pathology of BM fibrosis.

One of the most feared consequences of MPN for patients is clonal evolution to sAML. Approximately 20% of MF patients progress to sAML(9,41), which is associated with a dismal clinical prognosis and median survival of less than six months (41), largely due to the lack of targeted therapies. Genetic studies of paired chronic phase MF and sAML transformed samples from individual patients demonstrate that the clonal evolution of MF to sAML is driven by the acquisition of co-operating genetic mutations (9,29). Earlier identification of MF patients who are at-risk for sAML transformation may allow stratification for more aggressive therapies, bone marrow transplantation, or precision medicine approaches based on their mutation profile. However, the limit of detection of ~2% VAF for most clinical sequencing approaches combined with the fact that many of these assays only sample a portion of the genome prohibit this type of prediction. The PDX model presented here is able to identify rare leukemic subclones present in MF PBMC samples years prior to clinical diagnosis of sAML. Presumably, the engraftment of these subclones is favored in NSGS mice due to the selective pressure of xenotransplantation. The lack of such selective pressures in the chronic phase of MF patients means these clones may lay dormant for substantial periods of time. This is much like the case in therapy-related myeloid neoplasms where *TP53*-mutant subclones lay dormant until challenged with the selective pressure of chemotherapy (42). The engraftment of these clones in this PDX model facilitated identification of the relevant co-operating mutations responsible for disease progression in the patients, with these sAML-specific mutations far below the level of detection of exome sequencing at the time of sample collection. It could be argued that further advancements in technology will allow higher-resolution sequencing to identify these potentially transforming mutations at low abundance in MF patients during the chronic phase. However, error-corrected sequencing has shown that many potentially pathogenic mutations are acquired in people during age-related CH (43) and MDS (44), the vast majority of which are not functionally relevant without the right context. The power of this PDX system is that is able to identify variants with functional relevance by selection during the xenotransplantation process, with these variants being completely recovered in the eventual patient sAML. Thus, not only can this system prospectively identify rising sAML subclones in MF patients long before clinical presentation, it can serve as an avatar to test novel therapeutic approaches for individual patients based on their co-operating mutations.

As *JAK2*^V617F^, *CALR*^fs^ and *MPL*^W515^ mutations all converge on JAK/STAT signaling, the JAK1/2 inhibitor ruxolitinib is currently front line therapy for MF. Most patients experience a reduction in splenomegaly and an improved quality of life (45), but it does not eliminate the malignant clone and has little to no impact on BM fibrosis and overall survival (11). Moreover, 75% of MPN patients on ruxolitinib discontinue treatment within five years due to development of dose-limiting cytopenias and/or non-hematological toxicities (46). Therefore, there is a pressing need for new treatment modalities. However, pharmacological discovery in MPN has been hampered by the lack of appropriate models within which to evaluate the therapeutic effect on the key MF pathologies of BM fibrosis and MPN-initiating cell burden. We demonstrate here this PDX model can be used as a platform for clinical discovery for MF, recapitulating the patient experience with ruxolitinib of reduced splenomegaly without reducing MPN-initiating cell burden. Using this system, combination therapy of ruxolitinib with pan-PIM inhibition (currently in clinical trials for solid tumors) was able to effectively diminish MF patient cell engraftment and reduce splenomegaly. Most importantly, the combination of these agents was able to substantially reduce the burden of human *JAK2*^V617F^-mutant HSCs in the BM of these mice. While these results are encouraging, unfortunately this particular combination therapy has not proven feasible in MPN patients as myelosuppression has been observed with pan-PIM kinase inhibitors. It should be noted that while this PDX model may indeed be closer to the human disease, it is premature to imply it is fully predictive of human clinical outcomes with potential therapeutic agent(s). Nevertheless, our experiments serve as proof-of-principle that this system can serve as a platform for therapeutic screening. But in addition to constitutive JAK/STAT activation, other oncogenic signaling pathways including MAP kinase, PI3 kinase and NFκB pathways, are hyperactivated in MF patients (47–49). Combinatorial approaches with inhibitors of these pathways, many of which are FDA-approved for the treatment of other cancers, may yield therapeutic synergy with JAK/STAT inhibitors in MPN patients. These types of hypotheses can be readily tested using this system due to the lack of specialized techniques and materials, and this model should represent a generalizable tool which can be quickly adapted across labs in hematology research.

## METHODS

### Mice and transplantation

The Institutional Animal Care and Use Committee at Washington University approved all animal procedures. All mice used in this study were NOD-scid-*Il2rg*-null-3/GM/SF (NSGS; The Jackson Laboratory #013062). Human cells were transplanted into sublethally irradiated (200 rads) 6-8 week-old NSGS mice via X-ray guided (UltraFocus100, Faxitron) intra-tibial injections (right leg) in a volume of 30 μL into with 27-gauge U-100 insulin syringes (Easy Touch # 08496-2755-01). For intra-tibial injections, mice were anesthetized with an intraperitoneal injection of Ketamine/Xylazine mixture (2 mg/mouse; KetaVed, Vedco). Needle is inserted approximately 0.8 cm deep into the tibia and cells are gradually released into BM while the needle is gently removed. Supplementary Figure 1 details the number of CD34+ cells transplanted for individual patient samples.

### Human Samples

De-identified cord blood specimens were obtained from the St. Louis Cord Blood Bank. Peripheral blood samples were obtained from MPN patients with written consent according to a protocol approved by the Washington University Human Studies Committee (WU #01-1014) in accordance with the Declaration of Helsinki protocol. De-identified peripheral blood mononuclear cells (PBMCs) from MPN patients as well as from cord blood samples were isolated by Ficoll gradient extraction according to standard procedures. CD34+ cells were isolated using magnetic enrichment (Miltenyi Biotec #130-100-453) and cultured overnight in SFEMII media (StemCell Technologies #09605) supplemented with Pen-Strep (50 Units/mL), human stem cell factor (SCF; 50 ng/mL), human thrombopoietin (TPO; 50 ng/mL), and human FLT3L (50 ng/mL).

### Flow Cytometry

All antibody staining was performed in HBSS buffer (Corning #21021CV) containing Pen/Strep (100 Units/mL; Fisher Scientific #MT30002CI), HEPES (10uM; Life Technologies #15630080) and FBS (2%; Sigma #14009C). Briefly, bone marrow (BM) cells isolated from tibias, femurs, and iliac crests were combined for calculating total BM from each mouse. 1.0×10^8^ cells/mL were suspended in complete HBSS and incubated on ice for 20 min with the desired antibodies listed in the table below. All antibodies were used at 1:100 dilutions.

| **Epitope** | **Conjugate** | **Vendor** | **Product ID** |
| --- | --- | --- | --- |
| anti-mouse CD45 | BV605 | BioLegend | 103139 |
| anti-human CD45 | APC | BioLegend | 368512 |
| anti-human CD71 | APCy7 | BioLegend | 334110 |
| anti-human CD235a | PE | BioLegend | 349105 |
| anti-human CD3 | FITC | BioLegend | 317306 |
| anti-human CD19 | PECy7 | BioLegend | 302215 |
| anti-human CD33 | BV421 | BD Biosciences | 565949 |
| anti-human CD34 | PE | BioLegend | 343606 |
| anti-human CD90 | PECy7 | BioLegend | 328124 |
| anti-human CD45Ra | BV421 | BioLegend | 304130 |
| anti-human Lineage | APC | BioLegend | 348803 |
| anti-human CD45 | Biotin | BioLegend | 304004 |
| anti-human CD38 | FITC | Invitrogen | 11-0388-42 |
| anti-biotin Streptavidin | APCy7 | BD Biosciences | 554063 |

### Western Blot

10 μg of protein samples harvested from xenografted human cells were separated in pre-casted 4-15% gradient SDS gels (Biorad, #456-1084) and transferred to nitrocellulose membranes (Millipore #IPVH00010). Membranes were subsequently probed with antibodies to detect pSTAT3 (Cell signaling, #9145S), pSTAT5 (Cell signaling, #4322S) and β-ACTIN (Santa Cruz, SC-47778). Detection was performed using horseradish peroxidase conjugated secondary mouse or rabbit antibody and chemiluminescence HRP substrate (Millipore #WBKLS0100).

### Cytospins and Histopathology

For cytospins, 5.0×10^5^ cells were collected in 100μL PBS/2% FBS from single cell preparations of bone marrow, added to Shandon Single Cytofunnels (Thermo Scientific #1102548) clipped to glass slides (FisherScientific #12-550-20), and centrifuged using a Cytospin 3 centrifuge (Shandon) set at 500 rpm for 5 min at medium acceleration. After drying the slides for 45 mins at room temperature, cells were stained with the Hema 3 stat pack (Fisher Scientific #23-123869) and images captured with a Nikon Eclipse E200 microscope equipped with an Infinity 2 color camera (Lumenera) controlled by Infinity Capture software (Lumenera). For histopathology, mouse tibias and spleens were fixed in 10% neutral buffered formalin (Fisher Scientific #SF100-4) overnight at 4°C or 1 hr at room temperature. Spleens were hydrated as follows: 20% EtOH for 20 mins, 30% EtOH for 20 mins, 50% EtOH for 20 mins and 70% EtOH for 20mins. Immediately after overnight fixation, tibias were decalcified in 14% EDTA for 12 days. This is followed by series of hydration steps: 20% EtOH for 1hr, 30% EtOH for 1hr, 50% EtOH for 1hr and 70% EtOH for hr. Both tibias and spleens were rinsed with PBS and processed for paraffin embedding and sectioned at 5 μM. Both H&E and reticulin staining was performed by the Washington University Musculoskeletal Histology and Morphometry Core. Images were captured using an AxioObserver D1 inverted microscope (Zeiss, Thrownwood, NY) equipped with an Axiocam 503 color camera. Images were acquired with Plan-Apochromat objective at 63X, 1.4 NA objective using the ZEN 2 (blue edition) software.

### Single Cell RNA-seq

Cells were resuspended at 1,200 cells/uL in PBS + 0.04% BSA. cDNA was prepared after the GEM generation and barcoding, followed by the GEM-RT reaction and bead cleanup steps. Purified cDNA was amplified for 11-13 cycles before being cleaned up using SPRIselect beads. Samples were then run on a Bioanalyzer to determine the cDNA concentration. GEX libraries were prepared as recommended by the 10x Genomics Chromium Single Cell 3’ Reagent Kits v3 user guide with appropriate modifications to the PCR cycles based on the calculated cDNA concentration. For sample preparation on the 10x Genomics platform, the Chromium Single Cell 3’ GEM Library and Gel Bead Kit v3 (PN-1000075), Chromium Chip B Single Cell Kit (10x Genomics, PN-10000153), and Chromium Dual Index Kit TT Set A (PN-1000215) were used. The concentration of each library was determined through qPCR utilizing the KAPA library Quantification Kit according to the manufacturer’s protocol (KAPA Biosystems/Roche) to produce appropriate cluster counts Illumina NovaSeq6000 instrument. Normalized libraries were sequenced on a NovaSeq6000 S4 Flow Cell (Illumina) using the XP workflow and a 28×10×10×150 sequencing recipe. A median sequencing depth of 50,000 reads/cell was targeted for each library. scRNA-seq data were demultiplexed and aligned to the Genome Reference Consortium Human genome, CRChg38. The aligned data were then annotated and UMI-collapsed using Cellranger (v3.1.0, 10x Genomics). Cells with more than 10% mitochondrial gene expression and with top 5% unique feature counts were excluded. Principal component analysis was performed to reduce data using Seurat 3.0. Clusters were identified applying the dimensional reduction techniques (tSNE and UMAP) using the same package. Expression levels of conventional cell surface markers were examined to annotate clusters. Differentially expressed genes were identified within the cluster of interest. The fgsea package was employed to determine enriched signaling pathways.

### Exome Sequencing

Genomic DNA was extracted using the PureLink genomic DNA extraction kit (Invitrogen #K1820-02), and submitted for exome sequencing at the McDonnell Genome Institute at Washington University at an average coverage depth of 180x. Whole exome sequence data was aligned to reference sequence build GRCh38 using BWA-mem (1) version 0.7.10 (params: -t 8), then merged and deduplicated using picard version 1.113, (https://broadinstitute.github.io/picard/). Variants, including SNVs and indels, were detected using VarScan(2) version 2.4.2 (params: -- min-coverage 8 --min-var-freq 0.1 –min-reads 2). Combined SNVs and indels were annotated by DoCM (Database of Curated Mutations, params: --filter-docm-variants true), and further by Ensembl Variant Effect Predictor (VEP) of GRCh38 v95 (params: --coding-only false --everything --plugs [Downstream, Wildtype]) by providing gnomAD (The Genome Aggregation Database) and ClinVar VCF files. Variants were filtered by removing low quality variants (params: --min-base-quality 15, --min-mapping-quality 20), and removing sites that exceeded 0.1% population allele frequency in gnomAD projects. *CALR* frameshift mutations were manually called using BAM files in IGV. Primary sequence data is available at dbGAP.

### Droplet Digital PCR (ddPCR)

Genomic DNA was extracted as above. 20 ng of total genomic DNA was used to perform ddPCR. All primers and probes used for ddPCR were designed as per MIQE guidelines (50). Amplicon context sequences used to design these probes are as follows: EZH2 p.Y663H, hg19|chr7:148,507,406-148,507,528 and TP53 p.R248Q, hg19|chr17:7,577,477-7,577,599. Briefly, a PCR reaction mixture was prepared with 10 μL 2x Supermix for Probes without dUTP (Bio-Rad #1863023), 1 μL 20x target (FAM) and wild-type (HEX) primers/probe (Bio-Rad) and 9 uL of ddH2O. The PCR sample was partitioned into ~20,000 nanoliter-sized discrete droplets using the QX-100 Droplet Generator according to the manufacturer’s instruction. These droplets were then gently transferred into a 96-well plate (Bio-Rad #12001925), and ddPCR was performed using a thermocycler with following conditions: 95°C for 10 min, 40 cycles of 94°C for 30 s, 55°C for 60 s and 10 min at 98°C. The mutant allele frequencies were measured and calculated using Bio-Rad QX-100 droplet reader and QuantaSoft v1.7.4 software, respectively. Each plate contained at least one wild-type DNA control (Promega, #G152A).

### CRISPR/Cas9 and Targeted Deep Sequencing

CD34+ cells were cultured overnight in SFEMII media (Stemcell Technologies #09605) supplemented with Pen-Strep (50 Units/mL), human stem cell factor (SCF; 50 ng/mL), human thrombopoietin (TPO; 50 ng/mL), and human Flt3L (50 ng/mL). 12-24h post-culture, CD34+ cells were nucleofected with Cas9/ribonucleoprotein (IDT #1074181) complexed with gRNAs as previously described (39). Synthetic gRNAs sequences purchased from Synthego are as follows: *gPIM1.1:* GUGGCGUGCAGGUCGUUGCA and *gPIM1.2:* CUGGAGUCGCAGUACCAGGU. A gRNA targeting the *AAVS1* locus was used as a negative control: GGGGCCACUAGGGACAGGAU. 48 hrs post-nucleofection, approximately 100,000 cells were set aside to measure CRISPR/Cas9 targeting efficiency using PCR amplicon-based deep sequencing. The remaining cells were transplanted into sublethally irradiated (200 rads) NSGS mice via X-ray guided intra-tibial injection.

### Drug Treatments

CD34+ cells isolated from MF patients were transplanted into NSGS mice. Four-week posttransplant, xenografted animals were randomized into four groups to generate cohorts with equal levels of peripheral blood engraftment for treatment as follows: vehicle (5% dimethylacetamide/95% of 0.5% methylcellulose, 2x/day), INCB053914 (30 mg/kg, 2x/day), ruxolitinib, and INCB053914 plus ruxolitinib. Ruxolitinib was administered in chow formulation (2g ruxolitinib / 1kg chow; Incyte #INCB018424). Four-weeks post-treatment, effects were evaluated by flow cytometric and histological analysis as described above.

### Plasma Cytokine Analysis

Plasma serum was collected from 4-6 drops of peripheral blood obtained from NSGS mice via submandibular bleeding using Microtainer Collection Tubes (BD Microtainer, # 365967). Concentrations of 29 cytokines/chemokines were measured in 2-4 biological replicates using MILLIPLEX Map Human Cytokine/Chemokine Magnetic Bead Panel (HCYTMAG-60K-PX29, #Millipore) according to manufacturer’s protocol.

### Statistics

Student t-test, one-way, and two-way ANOVA’s were used for statistical comparisons where appropriate. Survival curves were analyzed using a Mantel-Cox logrank test. Significance is indicated using the following convention: *\*p* <0.05, \*\**p* <0.01, \*\*\**p* <0.001, \*\*\*\**p* <0.0001. All graphs represent mean ± S.E.M.

## DATA AVAILABILITY

Primary data is available under GEO accession number GSE160927.

## ACKNOWLEDGEMENTS

We thank all members of the Challen laboratory for critical discussions and support, particularly Jake Fairchild for animal husbandry and Samantha Burkart for laboratory oversight. We thank the Alvin J. Siteman Cancer Center at Washington University for use of the Siteman Flow Cytometry Core and Immunomonitoring Laboratory, supported in part by NCI Grant CA91842 and NIH WLC6313040077. We thank the Genome Technology Access Center and McDonnell Genome Institute at Washington University for genomic analysis, partially supported by NCI Grant CA91842 and by ICTS/CTSA NIH Grant UL1TR000448.

This work was supported by the National Institutes of Health (HL147978 to G.A.C.). H.C. was supported by an Edward P. Evans Foundation Young Investigator Award, an American Cancer Society Institutional Research Grant, the Leukemia Research Foundation, and the When Everyone Survives Foundation. G.A.C. is a scholar of the Leukemia and Lymphoma Society.

## AUTHOR CONTRIBUTIONS

Conceptualization: H.C., G.A.C. Data curation: H.C., R.C.Z.Z., T.L., S.T.O., G.A.C. Formal Analysis: H.C., R.C.Z.Z., T.L., G.A.C. Funding acquisition: H.C., G.A.C. Investigation: H.C., E.K., W.H., H.B., N.I., W.K.K. Methodology: H.C. Project administration: H.C., G.A.C. Resources: M.A., D.A.C.F., J.F., M.C.S., H.K.K., S.T.O. Software: R.C.Z.Z., T.L. Supervision: H.C., G.A.C. Visualization: H.C., R.C.Z.Z., G.A.C. Writing – original draft: H.C., G.A.C. Writing – review & editing: S.T.O., G.A.C.

## SUPPLEMENTARY FIGURE LEGENDS

**Supplementary Figure 1.**
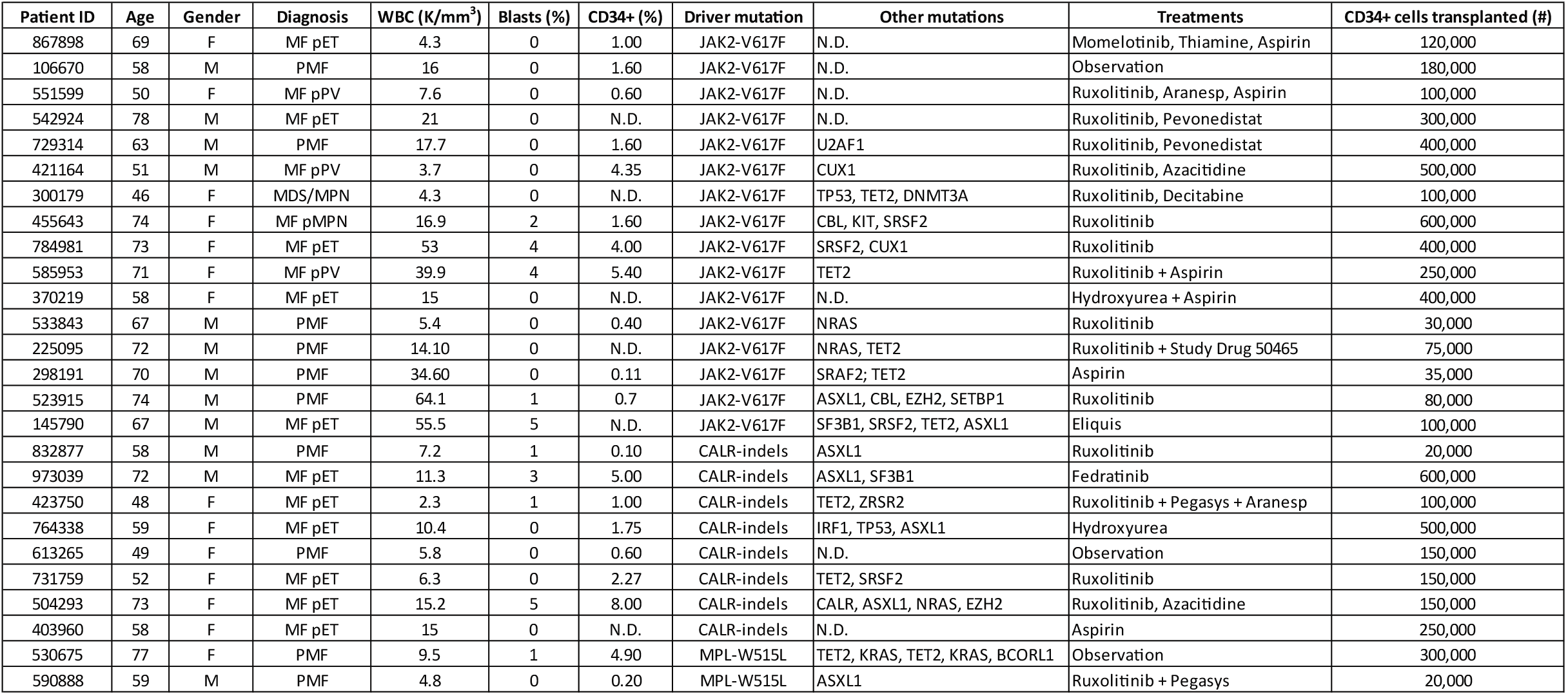
Characteristics of Myelofibrosis Patient Samples. Meta data for MF patient samples used in this study. Other mutations were extracted from clinical sequencing data, which were not comprehensive and typically only targeted to MF-specific mutations. N.D = not determined (not covered by clinical sequencing).

**Supplementary Figure 2.**
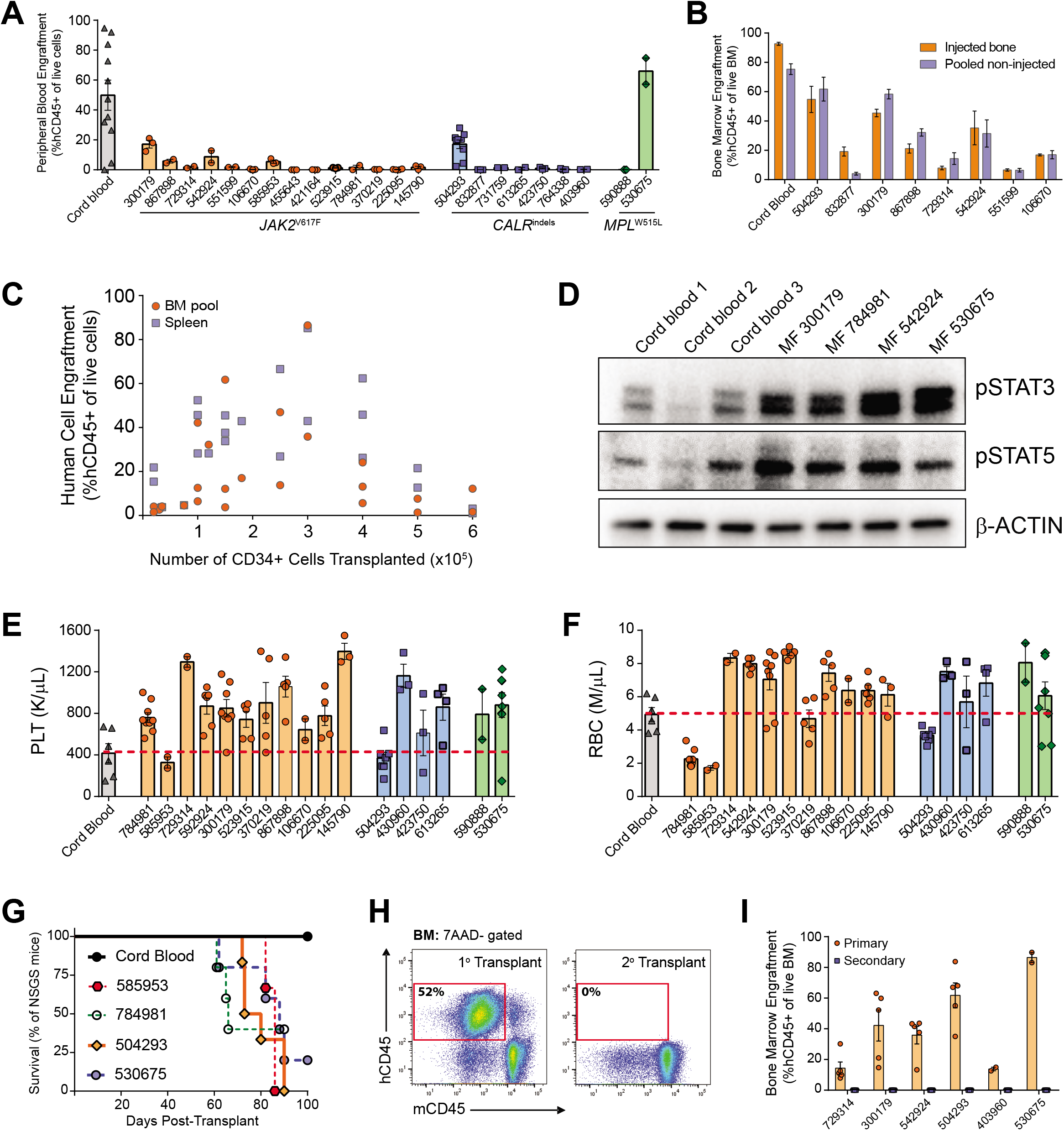
Credentialing a Humanized Animal Model of Myelofibrosis. **A**, Peripheral blood engraftment of human leukocytes 12-weeks post-transplant of MF patient samples. **B**, Engraftment levels of human cells in the BM of injected and pooled non-injected leg bones 12-weeks post-transplant. **C**, Correlation of BM and spleen engraftment of MF patient samples versus number of CD34+ cells injected. **D**, Western blot of hCD45+ cells purified from the BM of NSGS mice 12-weeks post-transplant showing activation of JAK/STAT signaling pathway in MF patient samples. Blood counts of platelets (**E**) and erythrocytes (**F**) 12-weeks posttransplant of indicated samples. Dashed red lines indicate average levels from xenotransplantation of cord blood CD34+ cells. **G**, Kaplan-Meier plot showing time to morbidity of NSGS mice transplanted with CD34+ cells from indicated MF patients compared to mice transplanted with cord blood CD34+ cells. **H**, Representative flow cytometric analysis of the BM of recipient mice 12-weeks post-primary and -secondary transplant MF patient sample 300179. **I**, 12-week engraftment of human cells in the BM of NSGS mice post-primary and -secondary transplant of individual MF patient samples. Error bars indicate mean ± S.E.M.

**Supplementary Figure 3.**
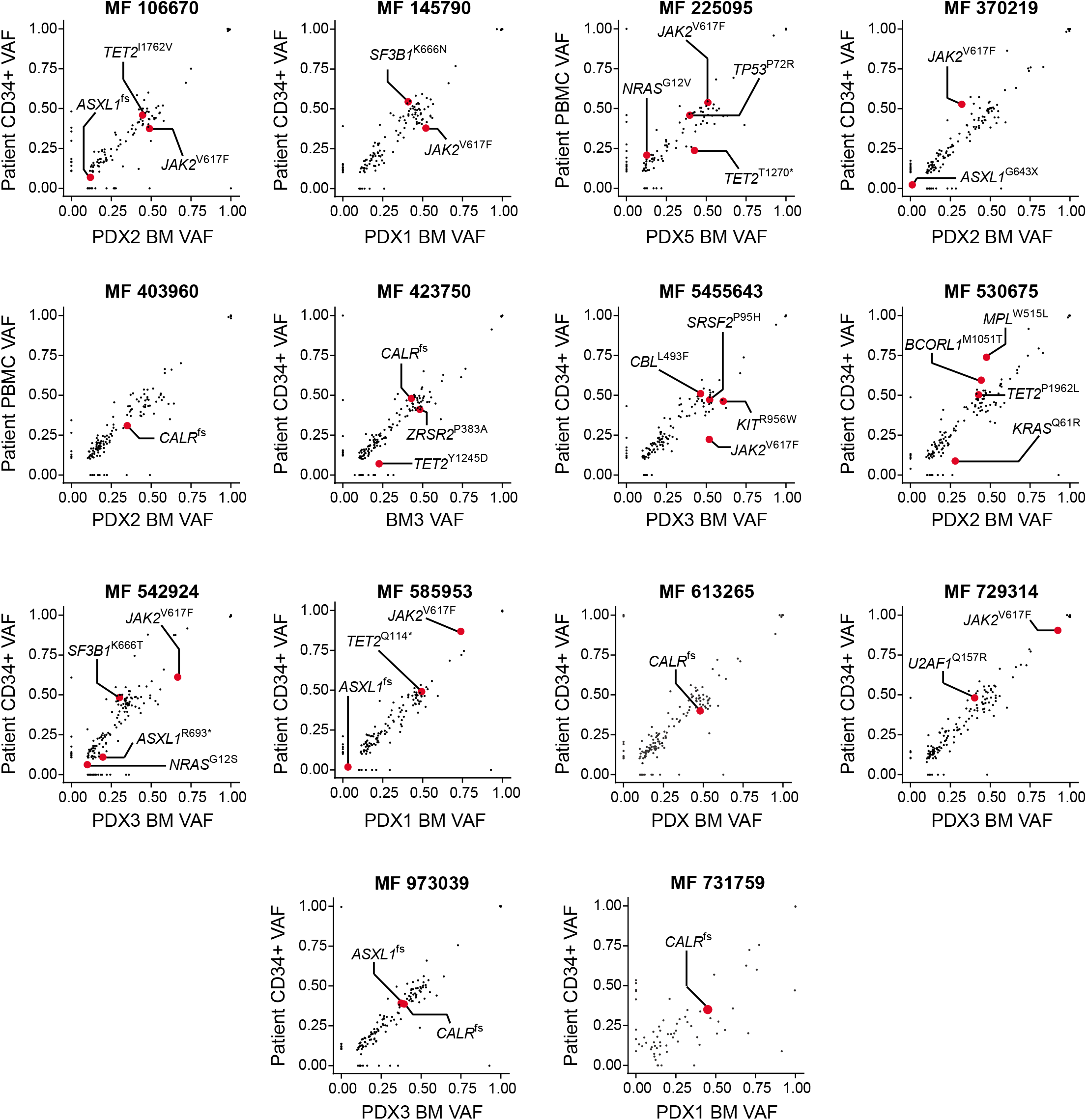
MF Patient Clonal Architecture is Maintained in PDX Models. Comparison of the variant allele fraction (VAF) of mutations in donor cells from MF patients versus hCD45+ cells isolated from BM of NSGS mice 12-weeks post-transplant.

**Supplementary Figure 4.**
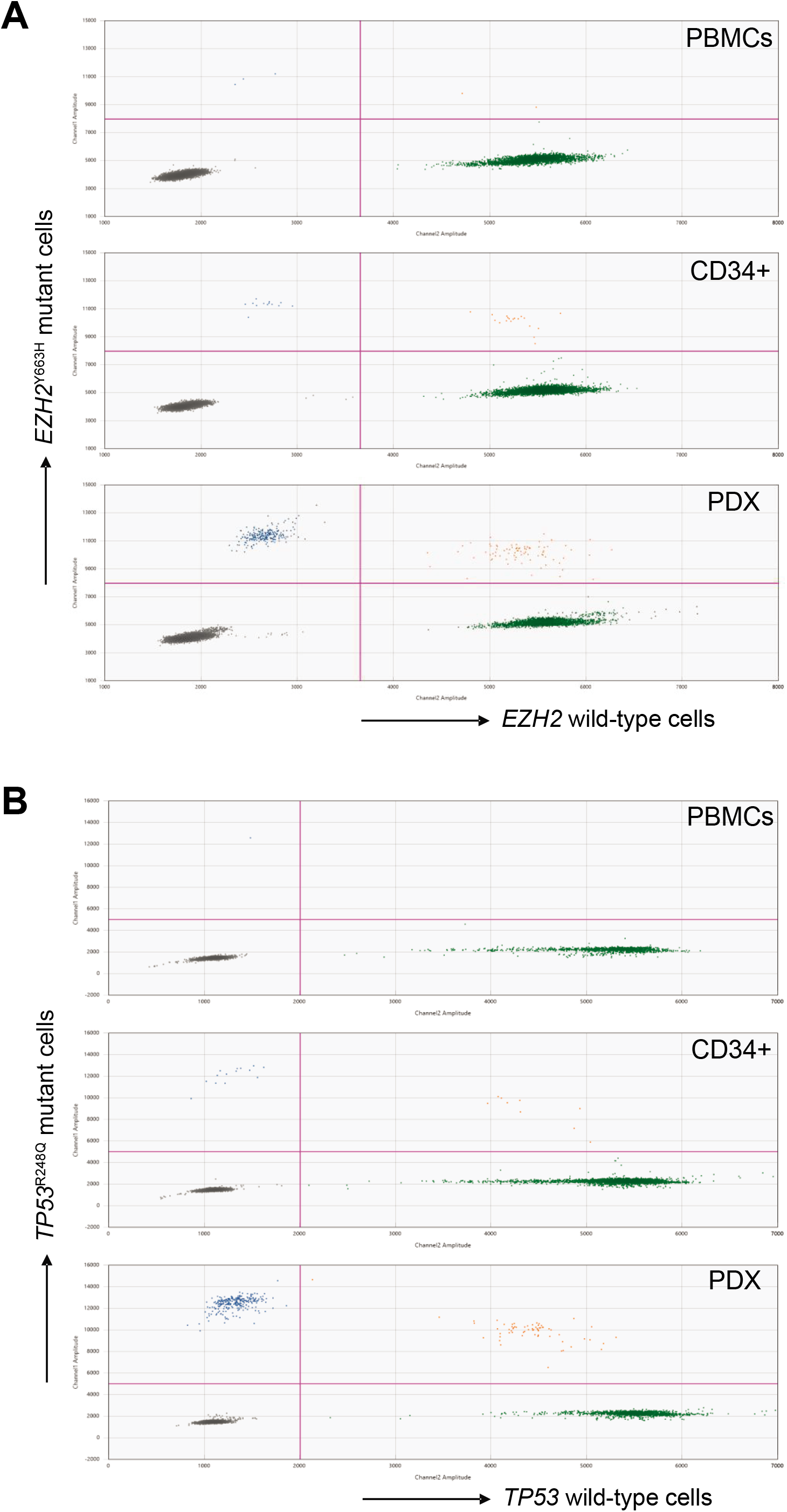
Rare Leukemic Subclones Are Present in MF Patients. **A**, ddPCR for *EZH2*^Y663H^ mutation in indicated samples of MF patient 504293. **B**, ddPCR for *TP53^R248Q^* mutation in indicated samples of MF patient 764338.

**Supplementary Figure 5.**
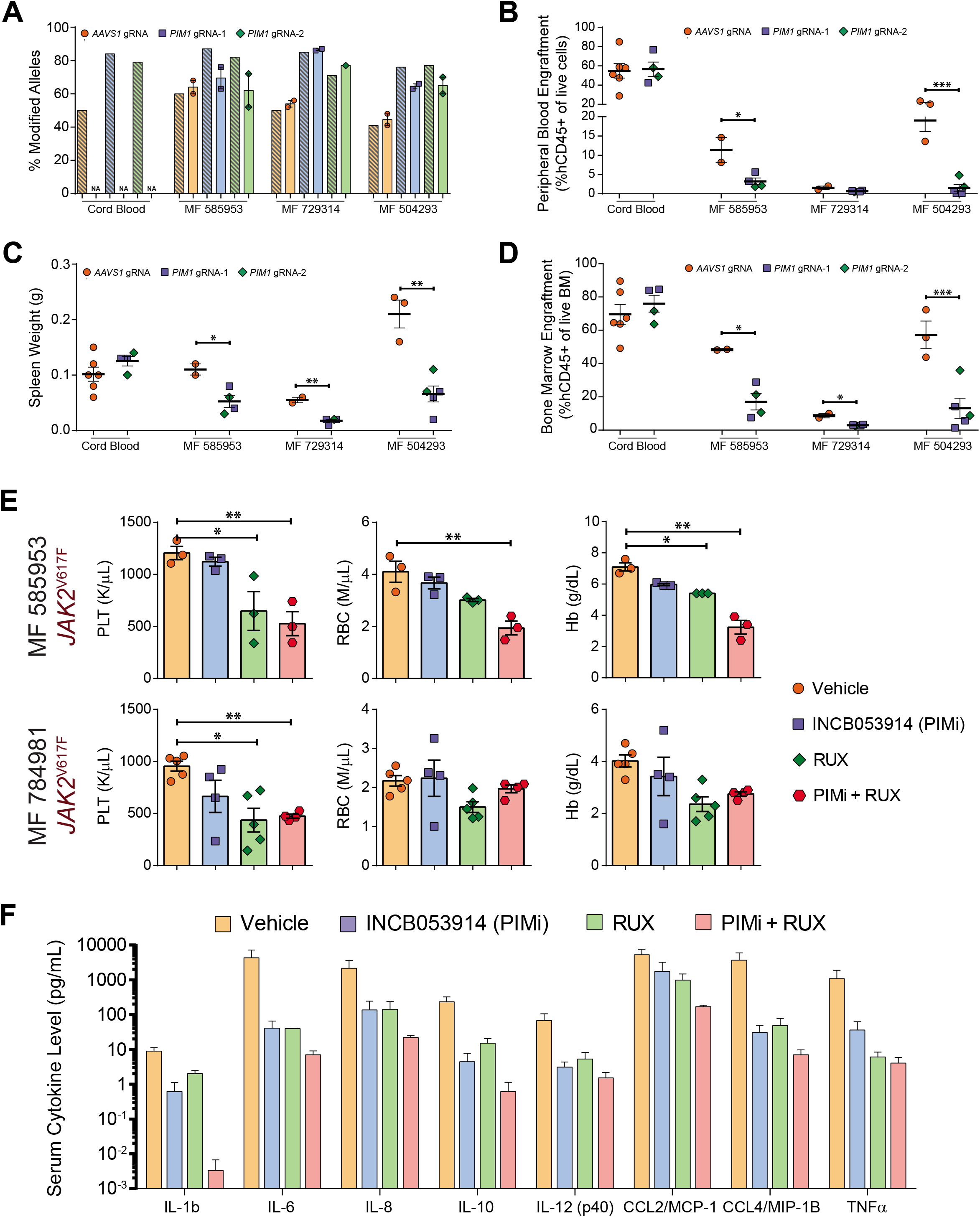
Genetic and Pharmacological Validation of Novel Therapeutic Targets in MF. **A**, Comparison of CRISPR/Cas9 targeting efficiency of indicated gRNAs pre- (shaded bars) and post-transplant (solid bars) in indicated patient specimens (N/A = sample not available). **B**, Peripheral blood engraftment of hCD45+ cells targeted with indicated gRNAs. **C**, Spleen weights of recipient mice transplanted with patient samples targeted with indicated gRNAs. **D**, BM engraftment of hCD45+ cells targeted with indicated gRNAs. **E**, Peripheral blood counts of NSGS mice transplanted with patient samples exposed to the indicated treatment arms. **F**, Serum concentrations of inflammatory cytokines in NSGS mice xenografted with MF patient samples and then exposed to indicated treatments. Serum samples were evaluated at the conclusion of therapy. Error bars indicate mean ± S.E.M. * *p*<0.05, ** *p*<0.01.

**Supplementary Table 1. Exome sequencing results.**

List of variants identified by exome sequencing in donor MF patient samples and xenografted human cells from NSGS mice 12-weeks post-transplant.

## REFERENCES

1. Campbell PJ, Green AR. The Myeloproliferative Disorders. New England Journal of Medicine 2006;355(23):2452–66.

2. Levine RL, Wadleigh M, Cools J, Ebert BL, Wernig G, Huntly BJ, et al. Activating mutation in the tyrosine kinase JAK2 in polycythemia vera, essential thrombocythemia, and myeloid metaplasia with myelofibrosis. Cancer Cell 2005;7(4):387–97.

3. James C, Ugo V, Le Couedic JP, Staerk J, Delhommeau F, Lacout C, et al. A unique clonal JAK2 mutation leading to constitutive signalling causes polycythaemia vera. Nature 2005;434(7037):1144–8.

4. Kralovics R, Passamonti F, Buser AS, Teo SS, Tiedt R, Passweg JR, et al. A gain-of-function mutation of JAK2 in myeloproliferative disorders. N Engl J Med 2005;352(17):1779–90.

5. Pikman Y, Lee BH, Mercher T, McDowell E, Ebert BL, Gozo M, et al. MPLW515L is a novel somatic activating mutation in myelofibrosis with myeloid metaplasia. PLoS Med 2006;3(7):e270.

6. Nangalia J, Massie CE, Baxter EJ, Nice FL, Gundem G, Wedge DC, et al. Somatic CALR mutations in myeloproliferative neoplasms with nonmutated JAK2. N Engl J Med 2013;369(25):2391–405.

7. Klampfl T, Gisslinger H, Harutyunyan AS, Nivarthi H, Rumi E, Milosevic JD, et al. Somatic mutations of calreticulin in myeloproliferative neoplasms. N Engl J Med 2013;369(25):2379–90.

8. Abdel-Wahab O. Genetics of the myeloproliferative neoplasms. Curr Opin Hematol 2011;18(2):117–23.

9. Abdel-Wahab O, Manshouri T, Patel J, Harris K, Yao J, Hedvat C, et al. Genetic analysis of transforming events that convert chronic myeloproliferative neoplasms to leukemias. Cancer Res 2010;70(2):447–52.

10. Verstovsek S, Kantarjian H, Mesa RA, Pardanani AD, Cortes-Franco J, Thomas DA, et al. Safety and Efficacy of INCB018424, a JAK1 and JAK2 Inhibitor, in Myelofibrosis. New England Journal of Medicine 2010;363(12):1117–27.

11. Tefferi A, Litzow MR, Pardanani A. Long-Term Outcome of Treatment with Ruxolitinib in Myelofibrosis. New England Journal of Medicine 2011;365(15):1455–7.

12. Mullally A, Lane SW, Ball B, Megerdichian C, Okabe R, Al-Shahrour F, et al. Physiological Jak2V617F expression causes a lethal myeloproliferative neoplasm with differential effects on hematopoietic stem and progenitor cells. Cancer Cell 2010;17(6):584–96.

13. Mullally A, Poveromo L, Schneider RK, Al-Shahrour F, Lane SW, Ebert BL. Distinct roles for long-term hematopoietic stem cells and erythroid precursor cells in a murine model of Jak2V617F-mediated polycythemia vera. Blood 2012;120(1):166–72.

14. Kent DG, Li J, Tanna H, Fink J, Kirschner K, Pask DC, et al. Self-renewal of single mouse hematopoietic stem cells is reduced by JAK2V617F without compromising progenitor cell expansion. PLoS Biol 2013;11(6):e1001576.

15. Klco JM, Spencer DH, Miller CA, Griffith M, Lamprecht TL, O’Laughlin M, et al. Functional heterogeneity of genetically defined subclones in acute myeloid leukemia. Cancer Cell 2014;25(3):379–92.

16. Nicolini FE, Cashman JD, Hogge DE, Humphries RK, Eaves CJ. NOD/SCID mice engineered to express human IL-3, GM-CSF and Steel factor constitutively mobilize engrafted human progenitors and compromise human stem cell regeneration. Leukemia 2004;18(2):341–7.

17. Song Y, Rongvaux A, Taylor A, Jiang T, Tebaldi T, Balasubramanian K, et al. A highly efficient and faithful MDS patient-derived xenotransplantation model for pre-clinical studies. Nat Commun 2019;10(1):366.

18. Lysenko V, Wildner-Verhey van Wijk N, Zimmermann K, Weller MC, Buhler M, Wildschut MHE, et al. Enhanced engraftment of human myelofibrosis stem and progenitor cells in MISTRG mice. Blood Adv 2020;4(11):2477–88.

19. Reinisch A, Thomas D, Corces MR, Zhang X, Gratzinger D, Hong WJ, et al. A humanized bone marrow ossicle xenotransplantation model enables improved engraftment of healthy and leukemic human hematopoietic cells. Nat Med 2016;22(7):812–21.

20. Celik H, Koh WK, Kramer AC, Ostrander EL, Mallaney C, Fisher DAC, et al. JARID2 Functions as a Tumor Suppressor in Myeloid Neoplasms by Repressing Self-Renewal in Hematopoietic Progenitor Cells. Cancer Cell 2018;34(5):741–56 e8.

21. Wunderlich M, Chou FS, Link KA, Mizukawa B, Perry RL, Carroll M, et al. AML xenograft efficiency is significantly improved in NOD/SCID-IL2RG mice constitutively expressing human SCF, GM-CSF and IL-3. Leukemia 2010;24(10):1785–8.

22. Saito Y, Ellegast JM, Rafiei A, Song Y, Kull D, Heikenwalder M, et al. Peripheral blood CD34(+) cells efficiently engraft human cytokine knock-in mice. Blood 2016;128(14):1829–33.

23. Yoshimi A, Balasis ME, Vedder A, Feldman K, Ma Y, Zhang H, et al. Robust patient-derived xenografts of MDS/MPN overlap syndromes capture the unique characteristics of CMML and JMML. Blood 2017;130(4):397–407.

24. Fleischman AG. Inflammation as a Driver of Clonal Evolution in Myeloproliferative Neoplasm. Mediators Inflamm 2015;2015:606819.

25. Jaiswal S, Fontanillas P, Flannick J, Manning A, Grauman PV, Mar BG, et al. Age-related clonal hematopoiesis associated with adverse outcomes. N Engl J Med 2014;371(26):2488–98.

26. Genovese G, Kahler AK, Handsaker RE, Lindberg J, Rose SA, Bakhoum SF, et al. Clonal hematopoiesis and blood-cancer risk inferred from blood DNA sequence. N Engl J Med 2014;371(26):2477–87.

27. Cordua S, Kjaer L, Skov V, Pallisgaard N, Hasselbalch HC, Ellervik C. Prevalence and phenotypes of JAK2 V617F and calreticulin mutations in a Danish general population. Blood 2019;134(5):469–79.

28. Matatall KA, Jeong MR, Chen SY, Sun DQ, Chen FJ, Mo QX, et al. Chronic Infection Depletes Hematopoietic Stem Cells through Stress-Induced Terminal Differentiation. Cell Rep 2016;17(10):2584–95.

29. Engle EK, Fisher DA, Miller CA, McLellan MD, Fulton RS, Moore DM, et al. Clonal evolution revealed by whole genome sequencing in a case of primary myelofibrosis transformed to secondary acute myeloid leukemia. Leukemia 2014.

30. Sabapathy K, Lane DP. Therapeutic targeting of p53: all mutants are equal, but some mutants are more equal than others. Nat Rev Clin Oncol 2018;15(1):13–30.

31. Rampal R, Ahn J, Abdel-Wahab O, Nahas M, Wang K, Lipson D, et al. Genomic and functional analysis of leukemic transformation of myeloproliferative neoplasms. Proc Natl Acad Sci U S A 2014;111(50):E5401–10.

32. Verbeek S, van Lohuizen M, van der Valk M, Domen J, Kraal G, Berns A. Mice bearing the E mu-myc and E mu-pim-1 transgenes develop pre-B-cell leukemia prenatally. Mol Cell Biol 1991;11(2):1176–9.

33. Horiuchi D, Camarda R, Zhou AY, Yau C, Momcilovic O, Balakrishnan S, et al. PIM1 kinase inhibition as a targeted therapy against triple-negative breast tumors with elevated MYC expression. Nat Med 2016;22(11):1321–9.

34. Brunen D, de Vries RC, Lieftink C, Beijersbergen RL, Bernards R. PIM Kinases Are a Potential Prognostic Biomarker and Therapeutic Target in Neuroblastoma. 2018;17(4):849–57.

35. Kapoor S, Natarajan K, Baldwin PR, Doshi KA, Lapidus RG, Mathias TJ, et al. Concurrent Inhibition of Pim and FLT3 Kinases Enhances Apoptosis of FLT3-ITD Acute Myeloid Leukemia Cells through Increased Mcl-1 Proteasomal Degradation. Clin Cancer Res 2018;24(1):234–47.

36. Padi SKR, Luevano LA, An N, Pandey R, Singh N, Song JH, et al. Targeting the PIM protein kinases for the treatment of a T-cell acute lymphoblastic leukemia subset. Oncotarget 2017;8(18):30199–216.

37. Huang SM, Wang A, Greco R, Li Z, Barberis C, Tabart M, et al. Combination of PIM and JAK2 inhibitors synergistically suppresses MPN cell proliferation and overcomes drug resistance. Oncotarget 2014;5(10):3362–74.

38. Norfo R, Zini R, Pennucci V, Bianchi E, Salati S, Guglielmelli P, et al. miRNA-mRNA integrative analysis in primary myelofibrosis CD34+ cells: role of miR-155/JARID2 axis in abnormal megakaryopoiesis. Blood 2014;124(13):e21–32.

39. Gundry MC, Brunetti L, Lin A, Mayle AE, Kitano A, Wagner D, et al. Highly Efficient Genome Editing of Murine and Human Hematopoietic Progenitor Cells by CRISPR/Cas9. Cell Rep 2016;17(5):1453–61.

40. Mazzacurati L, Collins RJ, Pandey G, Lambert-Showers QT, Amin NE, Zhang L, et al. The pan-PIM inhibitor INCB053914 displays potent synergy in combination with ruxolitinib in models of MPN. Blood Adv 2019;3(22):3503–14.

41. Mesa RA, Li CY, Ketterling RP, Schroeder GS, Knudson RA, Tefferi A. Leukemic transformation in myelofibrosis with myeloid metaplasia: a single-institution experience with 91 cases. Blood 2005;105(3):973–7.

42. Wong TN, Ramsingh G, Young AL, Miller CA, Touma W, Welch JS, et al. Role of TP53 mutations in the origin and evolution of therapy-related acute myeloid leukaemia. Nature 2015;518(7540):552–5.

43. Young AL, Challen GA, Birmann BM, Druley TE. Clonal haematopoiesis harbouring AML-associated mutations is ubiquitous in healthy adults. Nat Commun 2016;7:12484.

44. Duncavage EJ, Jacoby MA, Chang GS, Miller CA, Edwin N, Shao J, et al. Mutation Clearance after Transplantation for Myelodysplastic Syndrome. N Engl J Med 2018;379(11):1028–41.

45. Pardanani A, Tefferi A. How I treat myelofibrosis after failure of JAK inhibitors. Blood 2018;132(5):492–500.

46. Verstovsek S, Mesa RA, Gotlib J, Gupta V, DiPersio JF, Catalano JV, et al. Long-term treatment with ruxolitinib for patients with myelofibrosis: 5-year update from the randomized, double-blind, placebo-controlled, phase 3 COMFORT-I trial. J Hematol Oncol 2017;10(1):55.

47. Fisher DAC, Malkova O, Engle EK, Miner CA, Fulbright MC, Behbehani GK, et al. Mass cytometry analysis reveals hyperactive NF Kappa B signaling in myelofibrosis and secondary acute myeloid leukemia. Leukemia 2017;31(9):1962–74.

48. Kleppe M, Koche R, Zou L, van Galen P, Hill CE, Dong L, et al. Dual Targeting of Oncogenic Activation and Inflammatory Signaling Increases Therapeutic Efficacy in Myeloproliferative Neoplasms. Cancer Cell 2018;33(1):29–43 e7.

49. Fisher DAC, Miner CA, Engle EK, Hu H, Collins TB, Zhou A, et al. Cytokine production in myelofibrosis exhibits differential responsiveness to JAK-STAT, MAP kinase, and NFkappaB signaling. Leukemia 2019.

50. Hindson BJ, Ness KD, Masquelier DA, Belgrader P, Heredia NJ, Makarewicz AJ, et al. High-throughput droplet digital PCR system for absolute quantitation of DNA copy number. Anal Chem 2011;83(22):8604–10.

